## Supplemental Material for "Cataract induction in an arthropod reveals how lens crystallins contribute to the formation of ‘biological glass’"

### Supplementary materials

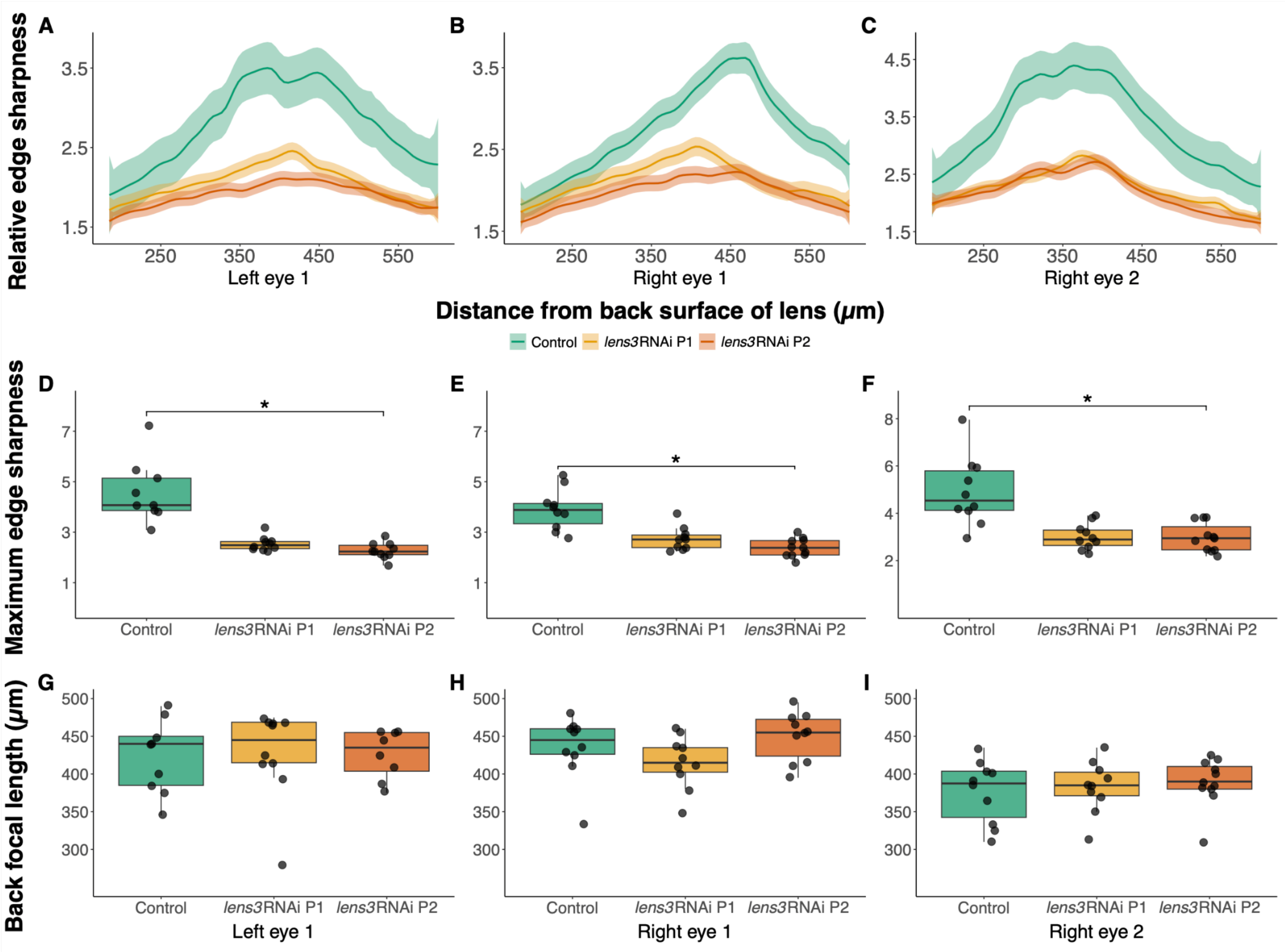


**Figure S1**. Assessment of lens optics through measurements of relative edge sharpness for left E1, right E1, and right E2. (A-C) Edge sharpness (LOESS) curves ($n_{control}$ = 9, $n_{lens3RNAi\ P1}$ = 10, $n_{lens3RNAi\ P2}$ = 11) show that the lenses of test individuals project blurry images compared to controls (shaded areas represent standard error). This is highlighted by a significant reduction in maximum edge sharpness (peak values extracted from individual curves) in lens3RNAi treated larvae (D-F; $n_{control}$ = 9, $n_{lens3RNAi\ P1}$ = 10, $n_{lens3RNAi\ P2}$ = 11; Kruskal-Wallis: d.f. = 2, $\chi^2_{LE1}$ = 19.7, $p_{LE1} < 0.01$; $\chi^2_{RE1}$ = 17.8, $p_{RE1} < 0.01$; $\chi^2_{RE2}$ = 14.8, $p_{RE2} < 0.01$;). (G-I) In contrast the back focal length, or the distance from the lens at which images are best focused (i.e., having highest edge sharpness), is not significantly different across groups (G-I; $n_{control}$ = 9, $n_{lens3RNAi\ P1}$ = 8, $n_{lens3RNAi\ P2}$ = 9; ANOVA: $F_{LE1;\ 2,25}$ = 0.13, $p_{LE1}$ = 0.87; $F_{RE1;\ 2,27}$ = 2.20, $p_{RE1}$ = 0.13; $F_{RE2;\ 2,28}$ = 0.32, $p_{RE2}$ = 0.72), suggesting that *lens3* knockdown does not cause detectable shifts in focal length.

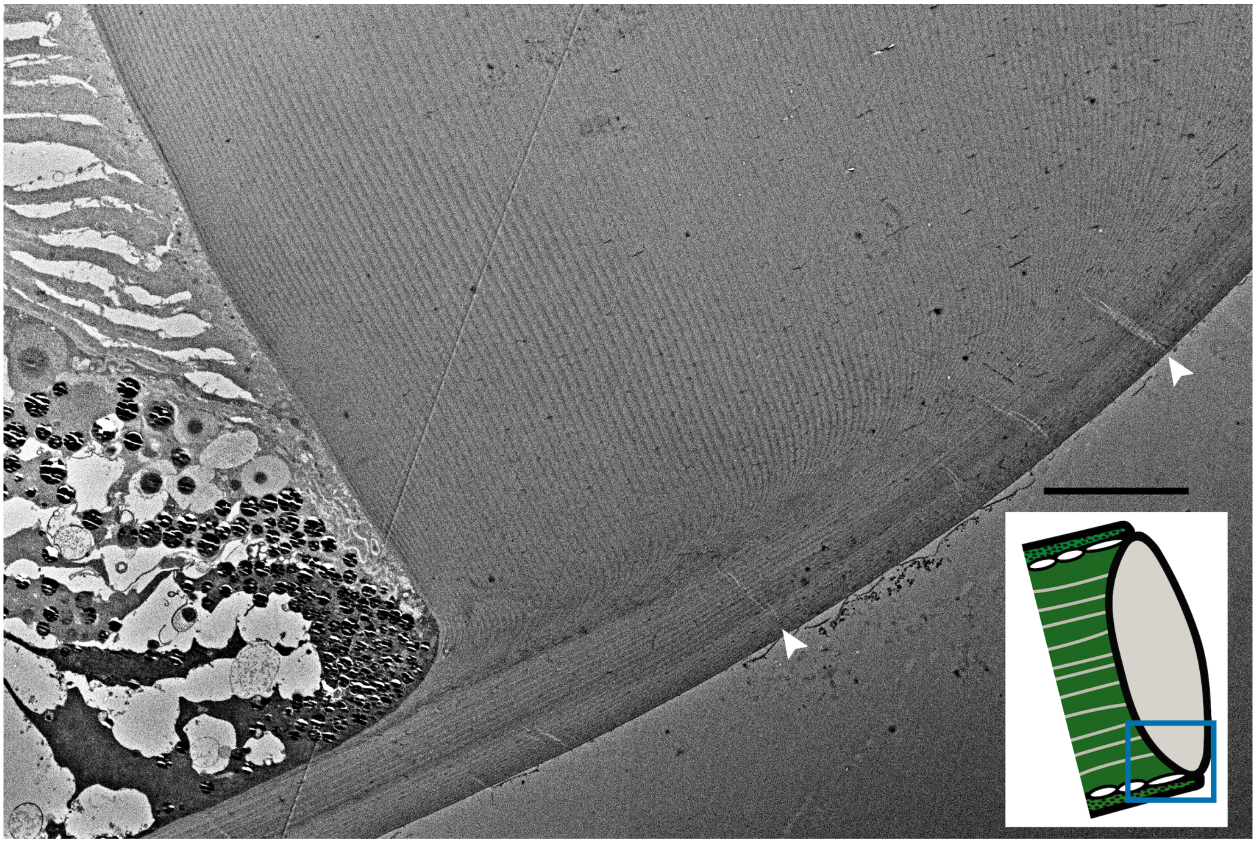

**Figure S2.** Opacities caused by reduced Lens3 levels are localised. In contrast to the proximal lens centre (Figure 5), the periphery and outer surface of *lens3*RNAi individuals show no defects, as exemplified in this TEM micrograph. Arrowheads point to pore canals typically found in arthropod lenses and cuticle [72,73]. Inset depicts the principal eye region that is visualised. Scale bar = 10µm.

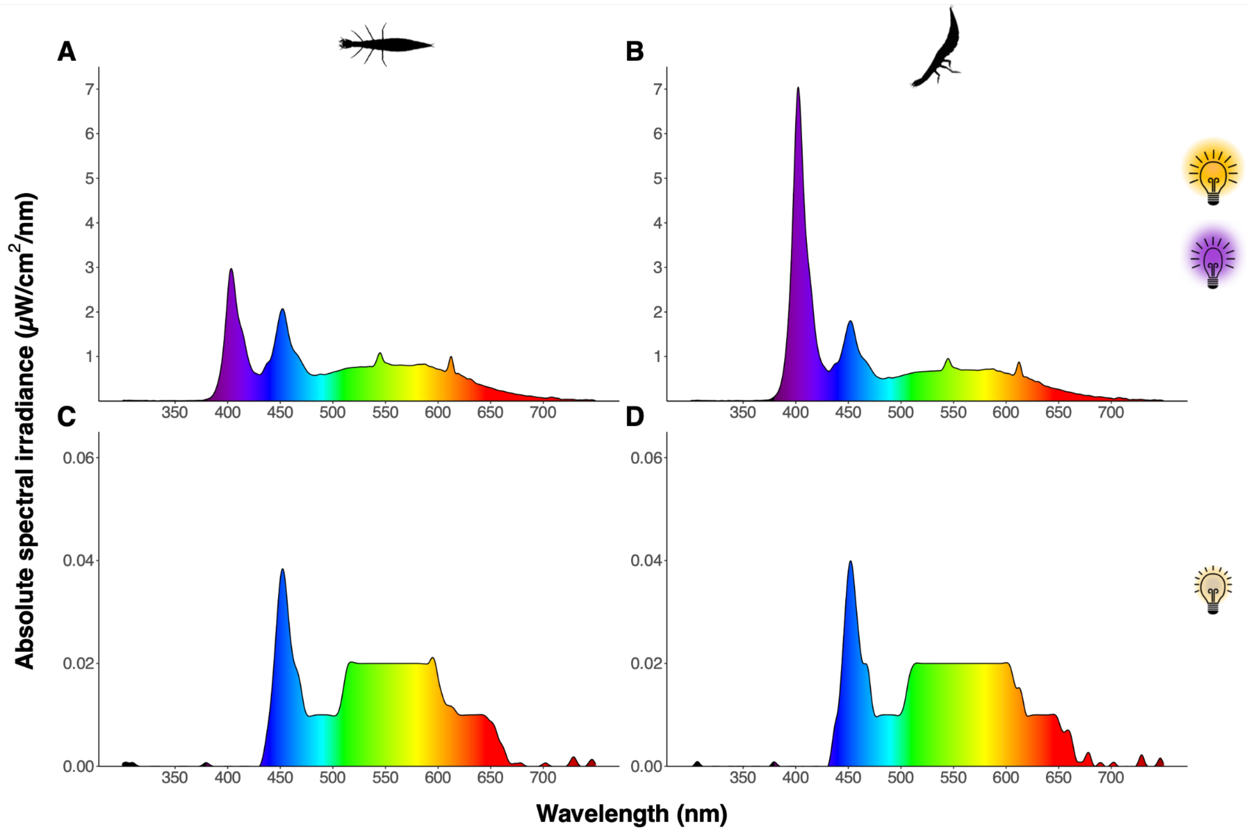


**Figure S3**. Spectral composition of lighting that was used in behavioural trials for assessment of larval hunting. Top row depicts horizontal (A) and vertical (B) arenas in bright white light supplemented with UV (horizontal arena = 6.27 x $10^{14}$ photons/$cm^2$/s; vertical arena = 7.03 x $10^{14}$ photons/$cm^2$/s). Bottom row depicts horizontal (C) and vertical (D) arenas in dim white light in the absence of UV supplementation (horizontal arena = 9.29 x $10^{12}$ photons/$cm^2$/s; vertical arena = 9.29 x $10^{12}$ photons/$cm^2$/s).

**Supplementary Table 1**. Sequence of T.marmoratus *lens3* gene identified through transcriptomics, and sequences of primers used for dsRNA synthesis and qPCR.

| Gene | | Obtained sequence (5' - 3') |
|---|---|---|
| *lens3* | | TTCCTCCAGCTCATTATGATTTTCAATATGGTGTAGATGAMMCAGATCACT<br>CACCGAGTCGATACAAATACGTTATACAGACATGTATTCCAAATTTGTCGTT<br>CTCGCCGCATTTGTGGCCATCGCTGCTGCCCAATACGGCCAATTTCAAGG<br>TCGTGAGGATCAACATGATGACACTTTTGCTCCAGCTCATTATGAATTTCAA<br>TATGGTGTAGATGACCCCAGCACTGGAGACAGAAAAACCCARCATGAAAC<br>CCGCCAAGGAGACGTTACCCAAGGAGAATACTCCGTAGTTGACCCTGATG<br>GCACCACCCGTACTGTCAAATACGCTGTCAACAGAAACTCTGGATTCCAA<br>GCTGAAGTCACCCGCTCCGGCCAAGCTCAGCACCCAACTCGTGATCAAC<br>TTAACCAAGGACCAGTAGCTGTGGCTCGCTCTCCAGTAGCTGTACATACC<br>TCAGTTGCTCCAGTATCAGTGCACCACGCCCCAATCGCCGTTGAACACGC<br>CCCAATTGCTGTCCACCCAACACCACTGTCTATCCACCCGACACCATTGG<br>CTGTCCAATCTACTCCAGTCGCTGTTCACCATACACAGATCCCAGTACAC<br>CCAACCCCAATAGTACACCACACCCCGATCCAAAACGGACCAATCGTTGA<br>TGCTTTCGGACATGGACAGCAAAACCAACGTGGATTTTTTAAATATAAATA<br>GGCCCAAGTAATTAAATTCCATATCATTCCATTTAACAAAAATAAAAATTATG<br>AAATAGTTTGTAGTTAGAAACCTCTAAATGTAAATAAATATTATTTACTTTAAA<br>ATGGTCAGTTATATTTTCATCCTGAAAATAGAAGAACCAACATGAATTAGAC |
| | **Probe 1 dsRNA amplicon primers with T7 linkers** | |
| | Forward | TAATACGACTCACTATAGGGTGGACAGCAAAACCA |
| | Reverse | TAATACGACTCACTATAGGGGTTGGTTCTTCTATT |
| | **Probe 2 dsRNA amplicon primers with T7 linkers** | |
| | Forward | TAATACGACTCACTATAGGGCGCTCTCCAGTAGCT |
| | Reverse | TAATACGACTCACTATAGGGGTTTTGCTGTCCATG |
| | **qPCR primers** | |
| | Forward | ATCTACTCCAGTCGCTGTTCAC |
| | Reverse | TTGCTGTCCATGTCCGAAAG |
| *RpL13A* | **qPCR primers** | |
| | Forward | ATGTCGTTCTTGCGCAAACG |
| | Reverse | AGCTTGCTTTCCACGTTCAG |
